# The Power of Citizen Science to Quantify Ecological Risks in Cities

**DOI:** 10.1101/2020.01.26.920124

**Authors:** Breanna J. Putman, Riley Williams, Enjie Li, Gregory B. Pauly

## Abstract

Urbanization is an extreme form of habitat modification that can alter ecological relationships among organisms, but these can be hard to study because much of the urban landscape is inaccessible private property. We show that citizen science can be a powerful tool to overcome this challenge. We used photo-vouchered observations submitted to the citizen science platform iNaturalist to assess predation and parasitism across urbanization gradients in a secretive yet widespread species, the Southern Alligator Lizard (*Elgaria multicarinata*), in Southern California, USA. From photographs, we quantified predation risk by assessing tail injuries and quantified parasitism rates by counting tick loads on lizards. We estimated urbanization intensity by determining percent impervious surface around each lizard observation. We found that tail injuries increased with age of the lizard and with urbanization, suggesting that urban areas are riskier habitats, likely because of elevated populations of predators such as outdoor cats. Conversely, parasitism decreased with urbanization likely due to a loss of mammalian hosts and anti-tick medications used on companion animals. Moreover, our citizen science approach allowed us to generate a large dataset on a secretive species extremely rapidly and at an immense spatial scale that facilitated quantitative measures of urbanization (e.g. percent impervious surface cover) as opposed to qualitative measures (e.g. urban vs rural). This study demonstrates that citizen science is allowing researchers to answer ecological questions that otherwise would go unanswered.

## Introduction

As the human population continues to grow and become more urban (1), animals increasingly have to survive and reproduce in modified habitats. Human-induced habitat modifications, like urbanization, shift ecological relationships including inter- and intra-specific competition and predator-prey and host-parasite interactions (2–4). Of specific interest are shifts in predation and parasitism as these have direct and indirect effects on species’ life history traits (5–7). If such relationships are altered so much that animals lack the phenotypic plasticity or evolutionary potential to respond, this could eliminate animals from urban habitats (8). Although extreme shifts in ecological relationships are likely to occur with increasing urbanization, there remain several barriers to studying them. First, much of the urban landscape is private property, and gaining access to conduct research can be logistically challenging if not impossible for larger studies. Second, and not unrelated to the first point, studying the ecology of organisms that are secretive or rare is logistically difficult because of the large temporal and/or spatial scale required to collect such an adequate dataset.

Citizen science (also called community science), which involves scientists partnering with members of the public to answer research questions, has the potential to fill these data gaps in urban ecology research. Platforms such as Zooniverse (https://www.zooniverse.org/), eBird (https://ebird.org/), and iNaturalist (https://www.inaturalist.org/) collate millions of observations of thousands of species worldwide. These large datasets of species occurrence records across large geographic scales have been used successfully to model species distributions, abundances, and overlap (9), to evaluate changes in phenology (10), and for biodiversity assessments (11). Most ecological studies using citizen science-generated species occurrence records use the spatial (i.e. latitude and longitude) and/or temporal (i.e. date and time) data associated with the observations. We expand on the uses of citizen science-generated data by using the spatial data in combination with the image content to quantify predation and parasitism, ecological interactions that affect fitness. Because citizen science observations cover the entire spectrum of urbanization and many of them come from private property, we can examine how these two key aspects of a species’ ecology change over a human-modified landscape.

There is conflicting evidence on the direction in which urbanization affects predation and parasitism. Predation could be lower in urban areas because of the human shield effect (3, 12) whereby predators are repelled by human presence, leading to relatively safe habitats for prey (13). Many urban animals, however, suffer serious predation by human companion animals such as owned or feral cats and dogs (14–16). As for parasitism, animals living in urban areas often have poorer body conditions (17, 18), are exposed to higher levels of environmental contaminants (19), and can experience higher population densities (20), all of which could enhance parasitic infections (21). Indeed, some urban populations of lizards (22), birds (23) and rodents (24) have higher levels of parasitic infections compared to non-urban populations. Nonetheless, the abundance and diversity of appropriate hosts/vectors is often reduced in urban environments, effectively lowering levels of parasitism in urban animal populations (25).

Here, we use photo-vouchered citizen science observations posted to the Reptiles and Amphibians of Southern California (RASCals) project (https://www.inaturalist.org/projects/rascals) to assess how the ecological risks of predation and parasitism scale with urbanization in a widespread, but secretive species, the Southern Alligator Lizard (*Elgaria multicarinata*). RASCals is hosted on iNaturalist and incorporates observations of herpetofauna from Southern California. Since its inception in 2013, over 49,000 observations have been added. Southern Alligator Lizards are the second most common species posted to RASCals, with over 5,500 observations as of 1 January 2020. Although commonly documented via citizen science, there are few published studies on the ecology of these lizards, likely due to the logistical difficulty in studying them. They are a solitary, secretive species found in both natural and urban habitats in the western United States and Mexico, and they generally avoid basking, preferring cooler temperatures and spending their time under logs, rocks, surface cover, vegetation, and other dark, moist microhabitats. Even experienced field ecologists are unlikely to see more than one a day in urban habitats and more than a handful per day in non-urban areas. However, as we demonstrate, crowdsourcing can generate numerous observations in a short period of time.

We measured predation risk by quantifying tail breaks in lizards from citizen science-generated photographs. Southern Alligator Lizards have long, semi-prehensile tails which they readily autotomize (self-amputate) as an escape tactic against predators. Tail autotomy is thought to be an indicator of predation intensity (26, 27); it can also occur through intraspecific aggression (28), but intraspecific aggression is not documented as resulting in tail autotomy in alligator lizards (29). The loss of the tail is not insignificant and has been shown to have serious fitness costs, including decreasing locomotor performance, increasing susceptibility to predation, lowering social status, increasing metabolism to replace lost tissue, and decreasing fecundity (27, 30). Thus, if tail breaks are positively affected by intensity of urbanization, then lizards living in urban environments might experience these fitness costs. Although several studies indicate animals have lower predation risk in urban areas, domestic or feral cats are common and kill an estimated 258 to 822 million reptiles each year in the United States (31). One study found that urban anole lizards in Puerto Rico had significantly more tail breaks than anoles living in natural areas, suggesting urban habitats are riskier than natural ones, but this is the only study of this kind (32). Based on this previous literature, iNaturalist observations of cats injuring and/or killing lizards (Fig. S1), and our own experiences with cats and lizards, we predicted that the probability of tail breaks would increase with urbanization.

In addition, we examined whether the tail ratio (relative length of unbroken tails) and break ratio (relative length of the original tail for lizards with tail breaks) vary with urbanization (Fig. 1A). Tail autotomy in lizards typically occurs in fracture planes within individual vertebra. Regrown tails lack vertebrae and instead have a stiff cartilaginous rod for support; thus, subsequent autotomy events tend to occur anterior to any prior break sites (33, 34). If the break ratio associates negatively with urbanization, this would suggest that tail breaks are occurring more frequently within individual lizards in urban environments (35). Variation in tail ratios along an urban gradient could indicate a shift toward altered morphologies due to selection, phenotypic plasticity, and/or spatial sorting.

**Figure 1.**
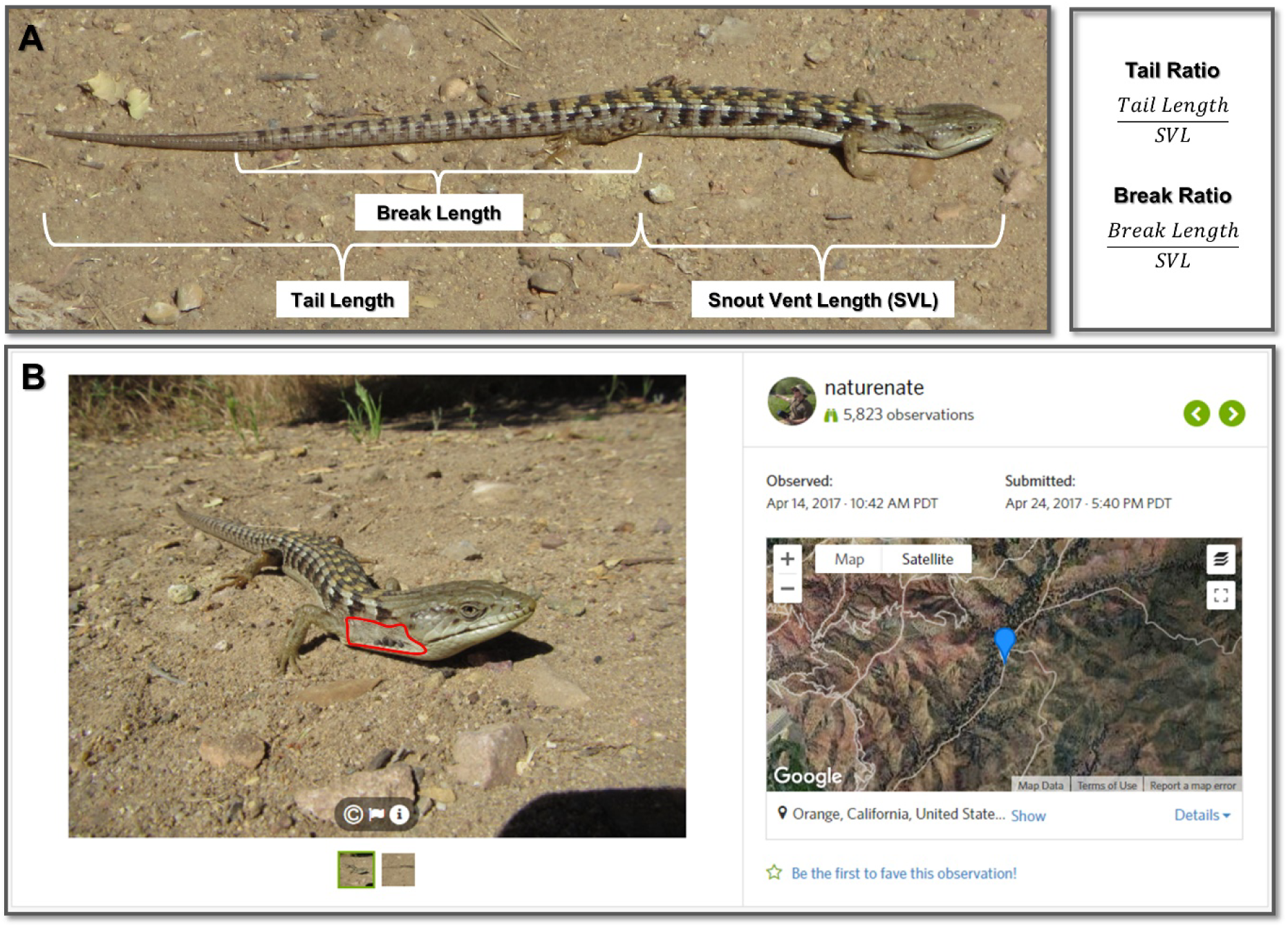
An observation of a Southern Alligator Lizard submitted to iNaturalist with two photographs, both of which were analyzed, showing the traits quantified in this study. (A) Body and tail measurements quantified to estimate predation risk. (B) The ear region (outlined in red) from which we estimated tick parasitism (with ticks present). The righthand side of the observation shows other data that were extracted including the date and time of observation, the date and time of submission, and the geographic locality. Photo and observation by Nathan Smith (iNaturalist 5945401; https://www.inaturalist.org/observations/5945401).

We measured parasitism by quantifying the prevalence and intensity of California Black-legged Tick (*Ixodes pacificus*) infection in lizards from citizen science observations (Fig. 1B). From photographs, we noted whether any ticks were present (ectoparasite prevalence) within the ear region and if so, we counted the number present (intensity of infection). Southern Alligator Lizards are a primary host of subadult ticks (36). Because the final hosts for adults of this tick species are large mammals, which are generally lower in abundance in urban areas (e.g. deer) or are treated with anti-tick medications (e.g. dogs and cats), we expected to see the prevalence and intensity of ectoparasite infections decrease with urbanization.

## Materials and Methods

Photo-vouchered observations of Southern Alligator Lizards (*Elgaria multicarinata*) were sourced from the Reptiles and Amphibians of Southern California (RASCals) project on iNaturalist (www.inaturalist.org/projects/rascals). We only included Research Grade observations that were submitted during a two-year period: October 2015 to September 2017. Research Grade means that observations have a photo voucher, locality information, date of the observation, and a community-supported taxonomic identification (Fig. 1). The locality information includes latitude and longitude as well as the locational accuracy. We only included observations with accuracy values less than 1km. For observations less than 100 m apart, we visually compared the size, color, and dorsal barring pattern of the lizards in the images to eliminate potential duplicates of the same individual. We quantified the mean percent impervious surface in a 100 m radius of each observation, based on the National Land Cover Database (NLCD)-2016 Percent Developed Imperviousness layer, and used this as a proxy for urbanization intensity.

Observations (N = 723) were included in the predation portion of the study if the entire tail was visible (including unbroken tails, broken tails that had not regrown, and regrown tails; Fig. 1A). We used the Segmented Line tool in ImageJ to quantify the number of pixels for the following: 1) snout-vent length (SVL) – from the tip of the snout to the location of the vent (cloaca), 2) tail length – from the vent to the posterior tip of the tail, and 3) break length – from the vent to the location of the break, if present (Fig. 1A). When a ventral view of the lizard was not available, which was the case with most observations, the vent was approximated to be 1–2 scale rows posterior to the hind limb insertion; this value was selected following examination of preserved *E. multicarinata* specimens at the Natural History Museum of Los Angeles County. We then calculated the tail ratio (tail length/SVL) for lizards with original unbroken tails and the break ratio (break length/SVL) for lizards that had experienced tail loss events.

Observations (N = 157) were included in the parasitism portion of the study if the entire ear region was visible (Fig. 1B) and individual scales in this region could be detected. This scale criterion was used because if these small scales could be detected, then individual nymphal ticks, which are a similar size, could also be counted if present. We chose to focus on the ear region because a preliminary examination of photographs and museum specimens revealed that this is the most common attachment area for ticks on *E. multicarinata*. We defined the ear region as the area with small scales on the side of the lizard between the ear opening and the forelimb insertion, above the ventral skin fold and below the larger, often keeled, scales of the dorsum (Fig. 1B). We excluded the ear opening as this was not consistently visible. We noted whether any California Black-legged Ticks (*Ixodes pacificus*) were present (ectoparasite prevalence) and if so, counted the number present (intensity of infection).

We categorized each lizard as either juvenile or adult based on dorsal pattern. Southern Alligator Lizards undergo an ontogenetic shift in color pattern with juveniles having a broad brown or tan stripe down their backs with no or minimal transverse dorsal bars; as they age, distinct dorsal bars appear. Juveniles were defined as lizards with incomplete or absent barring, while adults were described as individuals with complete barring (juvenile: *N* = 105, adult: *N* = 618).

From a subset of the observations we identified sex based on mating behaviors because males bite females on the neck, holding this position, before, during, and after mating (29). Based on submitted observations of mating behavior, we could determine the sex of a subset of individuals (female: *N* = 59, male: *N* = 75).

We examined the effects of urbanization (percent impervious surface) and age (juvenile or adult) on tail loss and ectoparasite prevalence using multiple logistic regressions. We looked for interactions between impervious surface and age and found none. We assessed whether the tail ratio and break ratio varied with age and urbanization using general linear models. Tail ratio and break ratio were log-transformed prior to analyses. We used observations with known sex (*N* = 134) to look for sex differences in tail loss, tail ratio, and break ratio and found none (all *P* > 0.05). We were unable to compare sex differences in ectoparasite presence because of small sample sizes (female: *N* = 1, male: *N* = 12). We did not include sex as a factor in further analyses. Furthermore, because only 20 individuals had tick infections (out of 157), we were unable to statistically assess the effect of urbanization on intensity of infection; thus, we report the results of logistic regression model on ectoparasite prevalence and descriptive statistics on intensity of infection. All analyses were run in R 3.6.1 (R Core Team, 2017).

## Results

Over a two-year period, 424 citizen scientists uploaded 1,688 Research Grade observations of Southern Alligator Lizards to the RASCals iNaturalist project. The dates of when animals were observed ranged from 8 May 2006 to 21 October 2017. Of the 1,688 observations, 723 met our criteria to be included in the predation study and 157 met the criteria to be included in the parasitism study. Importantly, the observations spanned a broad geographic range (Fig. S2) allowing us to quantify a continuous measure of urbanization intensity—percent impervious surface cover. This alligator lizard research was not advertised; we simply harvested observations from the RASCals project that fit the criteria. Observations were reviewed and selected for inclusion over a short time period (approximately 3 weeks) as time allowed, highlighting that citizen science platforms house tremendous opportunities for research.

The frequency of tail breaks increased with urbanization (N = 723, Z = 5.036, P < 0.001, Fig. 2A). For every 10% increase in impervious surface cover, the probability of tail loss increased by an average of 2.65 percentage points. In high density residential neighborhoods, the probability of tail loss was more than 20 percentage points higher than in a completely natural habitat (i.e. 0% impervious surface). Juveniles were less likely to have tail breaks than adults (N = 723, Z = −8.548, P < 0.001, Fig. 2A), likely because they have had less time exposed to risks.

**Figure 2.**
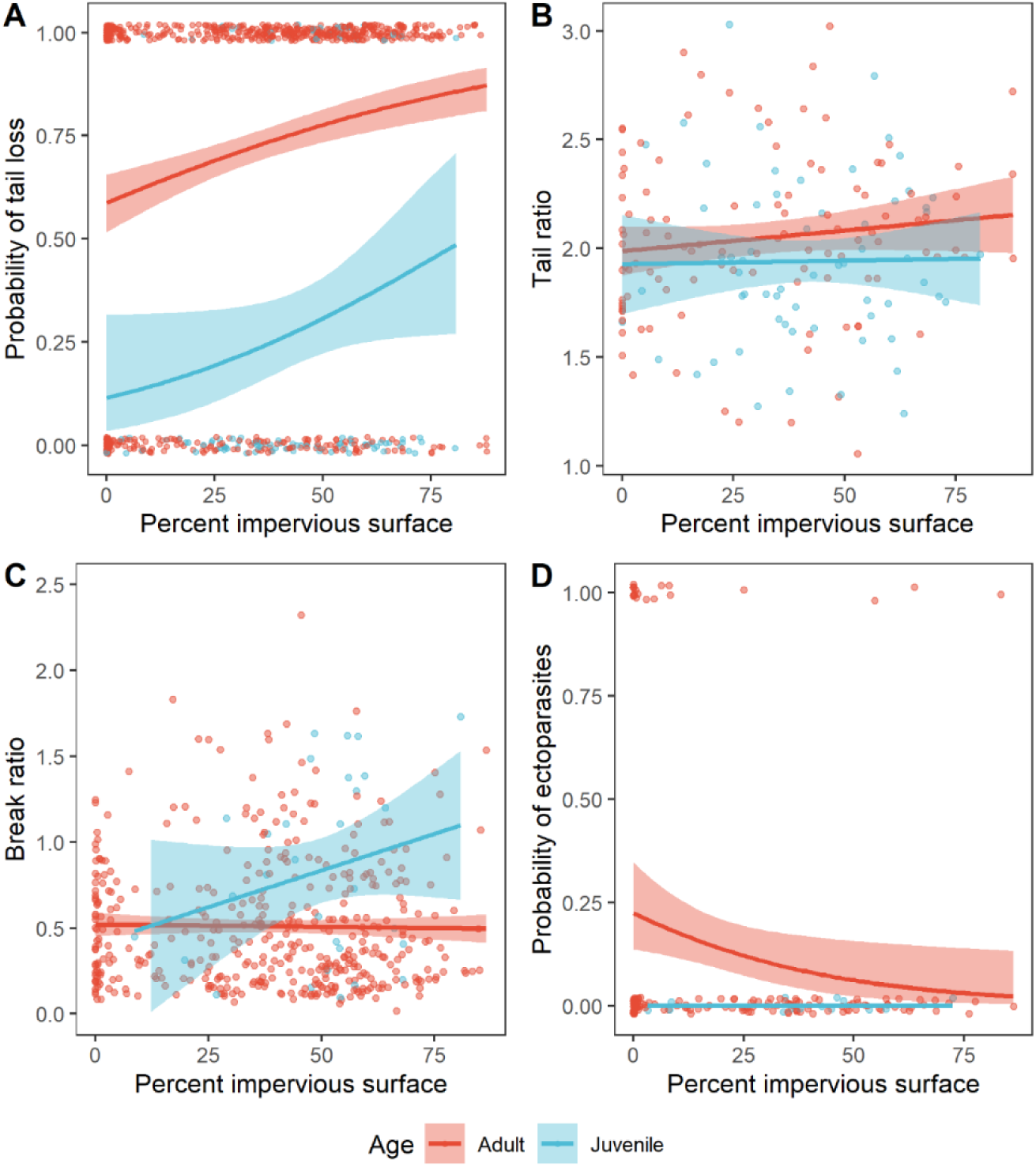
Effects of age and urbanization intensity on predation and parasitism in Southern Alligator Lizards. (A) Probability of tail loss (yes = 1, no = 0) (N = 723); (B) Tail ratio for lizards with unbroken tails (N = 171); (C) Break ratio for lizards that had experienced tail loss events (N = 455); (D) Probability of ectoparasitic infections (yes = 1, no = 0) (N = 157). Lines and standard errors generated from logistic regression models (A,D) or general linear models (B,C). For simplicity, the standard error has been removed from the “Juvenile” line in (D) because with no infected juveniles, the standard error is 1.

There was no effect of urbanization on the relative length of unbroken tails (i.e. tail ratio; N = 171, X^2^= 1.451, P = 0.228, Fig. 2B), but adults tended to have longer tails in relation to their body length compared to juveniles (X^2^= 3.746, P = 0.053). For alligator lizards that had suffered tail loss events, we did not find an effect of urbanization on the relative length of the original tail remaining (i.e. break ratio; N = 455, X^2^= 0.599, P = 0.439, Fig. 2C). Juvenile lizards with tail breaks, however, had significantly more of the original tail left in relation to their body size compared to adults with tail breaks (X^2^= 10.81, P = 0.001).

Ectoparasite prevalence decreased with urbanization (N = 157, Z = −2.25, P = 0.024, Fig. 2D). In urban habitats with 80% impervious surface cover, the probability of having ectoparasites was 2%, while this probability reached more than 20% in completely natural habitats (i.e. 0% impervious surface). It is notable that most infected lizards occurred in areas with less than 20% impervious surface, the cutoff for land classified as open space in the National Land Cover Database. Fifty percent of lizards with ticks in open space land had 3 or more ticks (maximum of 5), whereas all lizards found in more developed land classes had one or two ticks (Table S1). Age had no effect, although we did not find any ticks on juveniles in the dataset (Z = −0.010, P = 0.992).

## Discussion

We demonstrate that citizen science can be a powerful tool to answer modern ecological questions. Large datasets that span broad spatial scales were easily obtained in a relatively short time period. Importantly, we obtained data from areas of varying levels of urbanization, allowing us to examine changes in ecological risks across an urbanization gradient. Such data would have been unattainable via traditional research methods because (1) alligator lizards are secretive and generally have low detectability through traditional field surveys, and (2) most urban sites in Southern California are private property and therefore not easily accessible.

We obtained a large sample size of data on a secretive species more rapidly than those that use traditional field methods to study abundant and conspicuous species. For instance, although Tyler et al. (2016) collected 55–201 Puerto Rican crested anoles (*Anolis cristatellus*) at four sites to evaluate tail loss frequency in urban and rural populations, these data took three years to collect whereas we downloaded and analyzed over 700 photographs of lizards in less than a month. Furthermore, because citizen science-generated data cover a large geographic extent, we could evaluate the impacts of a continuous measure of urbanization on predation and parasitism. This is in contrast to many urban ecological studies that lump populations into urban versus non-urban categories (32, 37, 38) despite there being a great deal of variation across urbanized habitats (11).

Overall, we found both urbanization and age have significant effects on tails loss in alligator lizards. Juveniles were less likely to have tail breaks than adults, and probability of tail loss increased with higher levels of urbanization. Adult lizards living in the most urbanized areas have a 75–80% probability of having lost their tails, which is more than a 20-percentage point increase compared to adults living in natural sites. Thus, our data refute that these animals experience a human shield (i.e., humans shield them from natural predators). It is likely that for small-bodied, dispersal-limited organisms, such as most herpetofauna, cats are a major source of mortality in urban habitats (see 13). Tail breaks may also increase with urbanization because of a higher likelihood of encountering other risks such as vehicles, bicycles, and people. In support of this, one study found that Galapagos lava lizards (*Microlophus albemarlensis*) living near roads have a higher frequency of tail breaks than those farther from roads (39).

We did not find an effect of urbanization on the tail break ratio, the relative length of the original tail remaining for lizards that had experienced a tail break. This is an indirect measure for the number of tail loss events experienced by individual lizards as tail breaks are most likely to occur along vertebrae in the original tail. We found that adults had lower break ratios than juveniles, suggesting that lizards experience multiple tail breaks throughout their lives. Juveniles were also less likely to have tail breaks than adults. These two results corroborate past work which has shown that tail breaks accumulate with age (30). We also found that for lizards with their original tails, adults had longer tails relative to their body length compared to juveniles, suggesting positive allometric scaling of the tail occurs in this species.

As predicted, we found that ectoparasite prevalence decreased with urbanization. This is likely due to disruptions in the tick’s life cycle caused by urbanization. The final hosts for adult female ticks are medium to large mammals such as cervids, canids, and felids. Populations of these taxa are generally lower in abundance in urban areas than in natural ones. Domestic pets such as cats and dogs are also suitable final hosts, but humans often use anti-tick medications to reduce tick attachment on these animals. Thus, it appears that lizards are released from tick parasitism in increasingly urban areas. Urban lizards could therefore experience fitness benefits from reduced parasitism as costs of tick infestations include smaller home ranges, less movement, lower sprint speed, and lower endurance (40).

In sum, we demonstrate that citizen science platforms provide more than just spatial and temporal data of species occurrences. There is additional valuable information that can be gleaned from photographs including evidence of risk-associated trauma (tail breaks) and ectoparasite attachment (ticks). Our approach can be applied more broadly to study the ecology of other species in urban habitats or in other situations where crowdsourcing data collection could be more effective than using traditional approaches. The pace and scale of data collection through citizen science far exceeds traditional research methods and can help further our understanding of anthropogenic impacts on wildlife worldwide.

## Acknowledgments

We thank the many citizen scientists who uploaded observations to the RASCals project on iNaturalist and the developers of iNaturalist for building a platform that is revolutionizing biodiversity research. We thank the members of the Urban Nature Research Center (UNRC) and Community Science Office at the Natural History Museum of Los Angeles County for helpful feedback and discussion. This work was funded by a Postdoctoral Research Fellowship in Biology from the National Science Foundation (DBI-1611562 to B.J.P.) and by the Natural History Museum of Los Angeles County.

## Supplementary Material

**Fig. S1.**
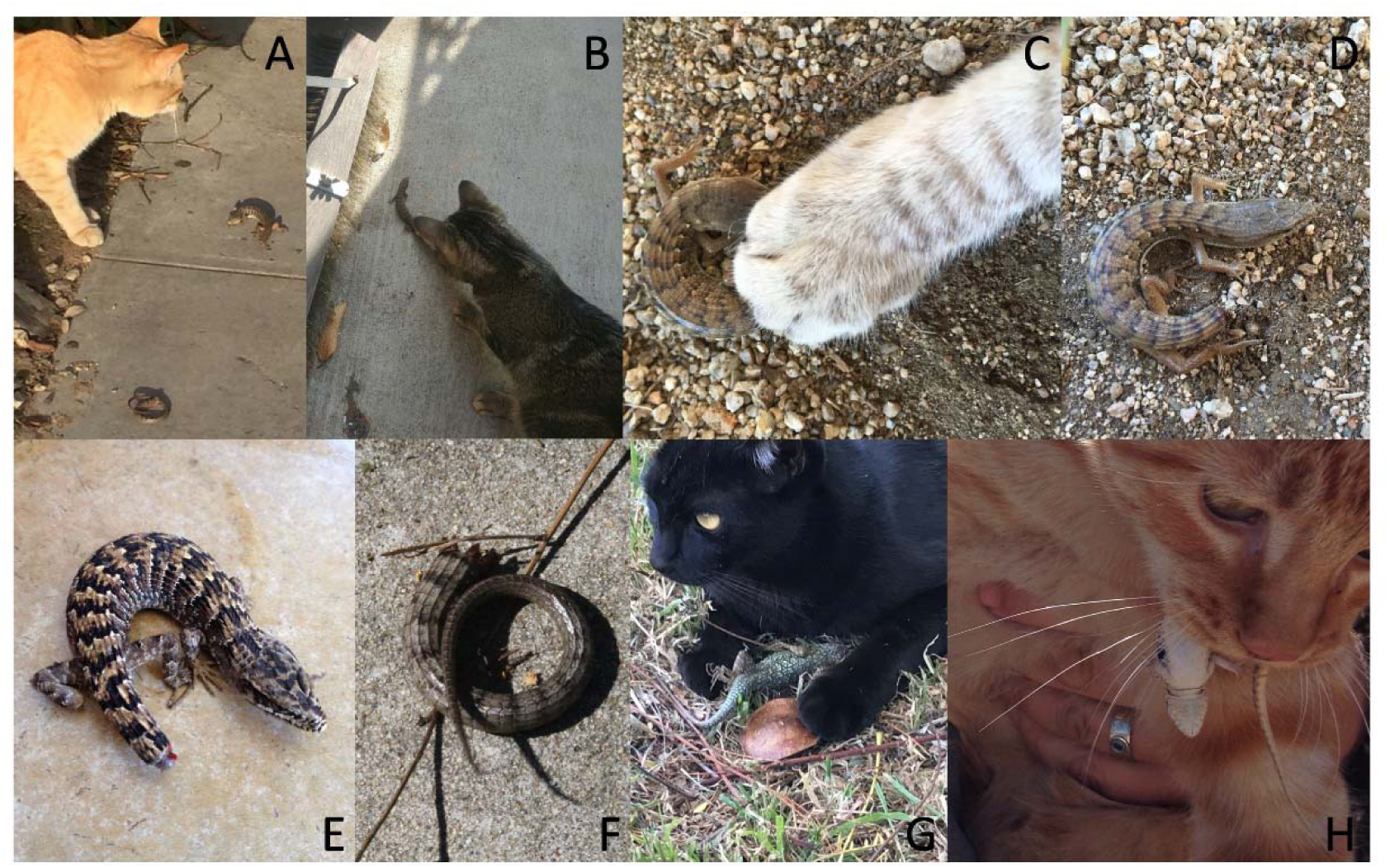
iNaturalist observations of lizards during or after interacting with cats in the Southern California study area. (A) Cat attacking a Southern Alligator Lizard (*Elgaria multicarinata*) that has autotomized its formerly complete, original tail; iNaturalist 2734344 by user ostevens. (B) Cat attacking a Southern Alligator Lizard that has autotomized its tail; iNaturalist 11288649 by user angus7. (C, D) Southern Alligator Lizard during and after an attack by a cat; iNaturalist 1733118 by Kristin Papoi and Violet Gibbs. (E) Southern Alligator Lizard with a tail injury after being caught by a cat; the fresh tissue at the tail tip is a re-growing tail following an autotomy event several weeks prior to this observation; iNaturalist 1447779 by Bruce Biesman-Simons. (F) Autotomized tail from a Southern Alligator Lizard on a neighborhood sidewalk following an attack by a cat; iNaturalist 2766323 by Patricia Simpson. (G) Western Fence Lizard (*Sceloporus occidentalis*) caught by a cat; iNaturalist 23869292 by user sharonn. (H) Side-blotched Lizard (*Uta stansburiana*) caught by a cat; iNaturalist 8274946 by Maiz Connolly, who noted that “as soon as the cat dropped the lizard, the lizard dropped its tail, which wriggled enough to keep the cat’s attention while it made its escape.”

**Fig. S2.**
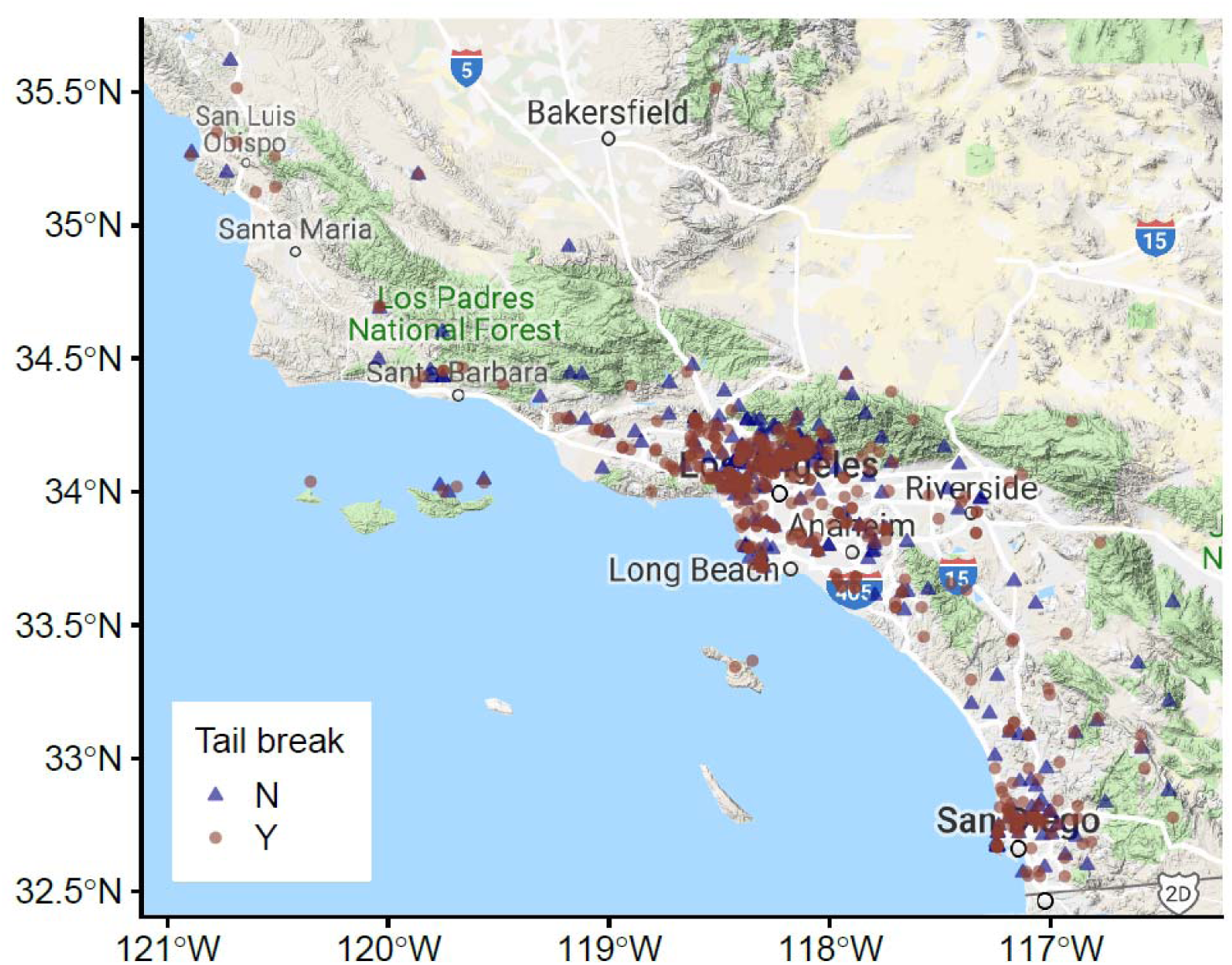
Map showing the geographic range of observations used in the predation portion of the study.

**Table S1.** Data associated with Southern Alligator Lizards that had tick infections.

| Age | Sex | Imperviousness | Number |
| --- | --- | --- | --- |
|  |  | (%) | of ticks |
| Adult | Unknown | 0.00 | 2 |
| Adult | Unknown | 0.00 | 5 |
| Adult | Unknown | 0.00 | 1 |
| Adult | Unknown | 0.00 | 1 |
| Adult | Unknown | 0.00 | 1 |
| Adult | Unknown | 0.00 | 1 |
| Adult | Unknown | 0.03 | 5 |
| Adult | Unknown | 0.29 | 5 |
| Adult | Unknown | 0.47 | 5 |
| Adult | Unknown | 0.62 | 3 |
| Adult | Unknown | 1.05 | 3 |
| Adult | Unknown | 2.97 | 2 |
| Adult | Unknown | 4.72 | 1 |
| Adult | Unknown | 6.32 | 3 |
| Adult | Unknown | 8.14 | 3 |
| Adult | Unknown | 8.41 | 1 |
| Adult | Unknown | 25.09 | 2 |
| Adult | Unknown | 54.80 | 2 |
| Adult | Unknown | 63.77 | 1 |
| Adult | Male | 83.31 | 1 |

